# Lethal Gene Drive Selects Inbreeding

**DOI:** 10.1101/046847

**Authors:** J J Bull

## Abstract

The use of ‘selfish’ gene drive systems to suppress or even extinguish populations has been proposed on theoretical grounds for almost half a century. Creating these genes has recently become possible with CRISPR technology. One seemingly feasible approach, originally proposed by Burt, is to create a homing endonuclease gene (HEG) that inserts into an essential gene, enabling heterozygote viability but causing homozygote lethality. With 100% segregation distortion in gametes, such genes can cause profound population suppression if resistance does not evolve. Here, population genetic models are used to consider the evolution of inbreeding (specifically selfing) as a possible response to a recessively lethal HEG with complete segregation distortion. Numerical analyses indicate a rich set of outcomes, but selfing often evolves in response to the HEG, with a corresponding partial restoration of mean fitness. Whether selfing does indeed evolve and its effect in restoring fitness depends heavily on the magnitude of inbreeding depression. Overall, these results point toward an underappreciated evolutionary response to block the harmful effects of a selfish gene. They raise the possibility that extreme population suppression may be more difficult to achieve than currently imagined.

## INTRODUCTION

The proposed use of selfish genes to suppress or extinguish populations is at least half a century old (Hickey and Craig 1966a,b; Hamilton 1967), but the feasibility of actually engineering selfish genes is new. There has thus been much excitement about the possibility of using these approaches to eradicate disease vectors, balanced by concerns about the possibility of unforeseen harm. Perhaps the most tangible approach is one outlined by Burt (2003), of creating a homing endonuclease gene (HEG) that inserts itself into an essential gene. Under the idealized assumptions of 100% segregation distortion in gametes of heterozygotes (germ line only), normal heterozygote viability and fertility but homozygote lethality, such a selfish gene is expected to evolve to such an extreme as to cause a 50% reduction in population fecundity if the segregation distortion is limited to one sex (Bruck 1957; Lewontin 1958). A segregation distortion that operates in both sexes can evolve to fixation and death of all progeny, ensuring extinction (Prout 1953; Lewontin 1958; Burt 2003).

The HEG need not work as completely as expected. Extreme levels of population suppression from the HEG are sensitive to even minor variations in parameter values (Deredec et al. 2008; Unckless et al. 2015). More importantly, an HEG that harms population fitness will select resistance mechanisms. Since HEGs target specific DNA sequences, the most obvious form of resistance to the HEG is a change in the target sequence so that the HEG will no longer duplicate itself in heterozygotes (Burt 2003). Resistance could also take the form of interfering with endonuclease expression or functionality. The problem of target sequence evolution has been countered with the suggestion of deploying multiple HEGs simultaneously (Burt 2003), but other resistance mechanisms would not obviously be thwarted by that approach.

Here I address another possible evolutionary mechanism that may interfere with the spread and long term maintenance of a lethal HEG: evolution of inbreeding. It is appreciated that inbreeding reduces the population impact of ‘lethal’ HEGs and other gene drive systems (Hamilton 1967; Burt 2003; Esvelt et al. 2014; Bull 2015). What is not clear is whether inbreeding is actually favored and how much it rescues mean fitness once a lethal HEG has invaded the population. Although a fixed level of inbreeding should reduce the incidence of the recessively lethal HEG, an allele that increases the level of inbreeding will itself suffer increased loss from any excess inviable progeny that it creates, perhaps selecting against inbreeding and even favoring increased outcrossing. It will in fact be shown here that inbreeding does evolve under some conditions, but the extent to which population fitness recovers depends heavily on the magnitude of inbreeding depression. Furthermore, the level of inbreeding that evolves is often not the level that would maximize population fitness were inbreeding imposed on the population.

The models analyzed here incorporate two major simplistic assumptions: the evolution of inbreeding is treated as the evolution of self-fertilization in an infinite population of simultaneous hermaphrodites with no spatial structure, and inbreeding depression is treated as a static quantity. Analysis of these models seems a reasonable first step in deciding whether the problem justifies inclusion of greater reality.

## THE MODELS

### Accommodating inbreeding as selfing

All models assume a population of diploid hermaphrodites capable of a mix of outcrossing and self-fertilization (selfing). Each individual produces a constant amount of sperm and of eggs. Eggs can be fertilized either by sperm chosen randomly from an out-cross pool or by self sperm, from the individual who produced the ova.

The models assume that selfed offspring have a fitness lower than that of outcrossed offspring (σ < 1), with inbreeding depression parameter

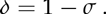

In this initial study, inbreeding depression is invariant throughout the evolutionary process. In real systems, inbreeding depression is often partially purged upon extended inbreeding, but allowing inbreeding depression to be static is a reasonable starting point and, if anything, provides a conservative measure of the vulnerability of gene drive systems to be suppressed by inbreeding (see Discussion).

### Genotypes and phenotypes

The models assume two unlinked loci, each with two alleles (*A/a, D /d*). An individual’s level of selfing is controlled by its genotype at the *A/a* locus independently of the genotype at the other locus (Table 1). The D */ d* locus affects viability and experiences gametic drive. Specifically D is a recessive lethal that enjoys complete segregation distortion in heterozygotes – *Dd* produces all D gametes in spermatogenesis and (in some models) also in ovogenesis. For most analyses, *Dd* is considered to be of normal viability and fertility (the drive does not operate in somatic tissues so those remain heterozygous and function normally); some analyses instead impart a slight fitness disadvantage to the heterozygote. The DD genotype of both sexes dies at conception, so viable genotypes at this locus are limited to dd and Dd.

**Table 1.** Genotype control of selfing rate

| Genotype |  | Proportion ova selfed |
| --- | --- | --- |
| aadd | aaDd | $s_0 = 0$ |
| Aadd | AaDd | $s_1$ |
| AAdd | AADd | $s_2$ |

### Modifications of reproductive osutput

The net reproductive output of a genotype that produces selfed offspring or experiences zygote loss can be modeled in different ways, each of which may be observed in nature. For example, there are two extremes that span the possibilities for the impact of selfing on sperm contributed to the outcrossed pool (Porcher and Lande 2005a; Harder et al. 2008). At one extreme, sperm may be reduced in proportion to the fraction of ova selfed (sperm are ‘discounted’). Alternatively, the outcross contribution may be unaffected (sperm not discounted) because few sperm are actually used for selfing compared to the huge number released for outcrossing. Both cases will be evaluated here.

In species with post-zygotic investment in offspring, reproductive compensation may also operate for lost zygotes, whereby the total number of viable offspring released from the maternal parent is larger than expected from the fraction of viable zygotes (Porcher and Lande 2005b; Harder *et al.* 2008). Here, the models are limited to maternal offspring production in direct proportion to viable zygotes.

### Four models

The foregoing sections identified two important biological variations that affect the evolutionary dynamics:

a. drive operates just in male gamete production or in both male and female gamete production
b. the contribution of sperm to the outcrossed pool is discounted in proportion to the fraction of ova selfed, or alternatively, selfing has no effect on the contribution to the outcross pool.

By considering these variables in all combinations, there are four models.

Interest is in whether selfing evolves specifically in response to the presence of *D* and the load from *DD* inviability. Appropriate analyses are thus limited to parameter values in which selfing would not evolve if *D* was absent. It is now well known that the evolution of selfing is sensitive to both the magnitude of inbreeding depression and whether selfed sperm are discounted (Lloyd 1979; Lande and Schemske 1985; Harder et al. 2008). When selfed sperm are not discounted from the outcross pool, there is an extra benefit to male function. Consequently, selfing is intrinsically beneficial at low values of inbreeding depression and is favored until inbreeding depression exceeds 0.5; the appropriate range for our problem is thus5 *δ* = 0.5. If instead selfed sperm are discounted, selfing is favored only if inbred offspring are more fit than outcrossed offspring, so the appropriate range for our problem includes even small values of inbreeding depression,*δ* >0.

The models will be analyzed for gene frequency evolution and mean fitness in the population (measured as viable offspring). Applied interest in these evolutionary processes is primarily for population control and the potential for extinction (Burt 2003,2014; Gould *et al.* 2008; Gould 2008). Yet, as has long been realized in applications of the sterile insect technique (used to suppress target pest populations), the impact of a particular intervention on adult population size depends heavily on ecology, which is often species-specific. Thus the analyses here omit any translation between mean fitness and population suppression.

## RESULTS

### Expectations

The evolution of gene drive has different consequences depending on whether drive operates in one sex (males, here) or both sexes.

A drive allele with 100% distortion in sperm and ova of the same *Dd* heterozygote can evolve to the extreme that all offspring in the population are inviable DD (Prout 1953). This outcome ensures population extinction. Drive operating only in one sex can only evolve to the point that half the offspring are DD, half are *Dd* (Bruck 1957; Lewontin 1958) with a 50% reduction in population fitness (fecundity). A minimal expectation is thus that selfing should be favored for higher levels of inbreeding depression when drive operates in both sexes than when it operates just in one sex. For example, one might anticipate that selfing would not evolve in response to male-drive when inbreeding depression exceeds 0.5 *(σ* < 0.5) because mean fitness in the absence of selfing is 0.5.

### Broad patterns in the joint evolution of selfing and gene drive

The four models were analyzed for a variety of initial conditions (Fig. 1). For each model, two different inbreeding depression values (δ) were chosen within the feasible range, and trials were conducted with different initial genotype frequencies and for different combinations of selfing rates (s1, s2). In each panel of Fig. 1, a solid blue line gives mean fitness in the absence of the HEG across different levels of inbreeding, specified on the horizontal axis. The line has slope —*δ*. A dashed orange curve shows mean fitness in the presence of the HEG for a fixed, uniform level of inbreeding. As selfing rate increases from 0, the yellow dashed curve rises until it intersects the blue line and then declines, coinciding with the blue curve. The rise in the yellow curve is from the reduced impact of the HEG on mean fitness under inbreeding. The HEG is lost for all selfing rates at which the yellow and blue curves coincide, so the decline over that segment is due to the loss in fitness from increased exposure to inbreeding depression.

Each panel includes several black triangles. These triangles are selected outcomes of evolution at the selfing locus, chosen from trials that collectively tested 120 different (s1, s2) values. The figures show only the outcomes at or near the highest evolved mean fitnesses observed across the trials. (The triangles exclude outcomes in which *D* was lost except those where *D* was lost because complete selfing evolved; see ‘Paradoxical’ section below.)

Fig. 2. illustrates the maximum fitness versus inbreeding observed from the triangles in each of the 8 panels of F(Fig. 1. Each of the 8 points is coded to indicate one-sex drive versus two-sex drive (1 versus 2) and whether sperm are discounted (triangles) or not (circles).

From the two figures, several points are noteworthy.

1. Given appropriate selfing rates for the *Aa* and A A genotypes, selfing evolved and increased mean fitness at least slightly above the fitness evolved with the HEG alone. Evolved fitness was equal to the fitness of selfed offspring *(σ)* for the sperm discounting models but somewhat above *σ* for the models with no sperm discounting.
2. Mean fitness did not necessarily attain the maximum that could be obtained if selfing was imposed on the population.
3. In the male-drive models, selfing never evolved if the fitness of selfed offspring *σ* was too low. Invasion analyses indicated that *σ* needed to exceed 0.5 in the sperm discounted model and 0.25 in models where sperm were not discounted.
4. The drive allele (D) was lost only when a selfing rate of 1 could evolve (see below for some exceptions, however).

Given that DD is lethal, the assumption that *Dd* is fully viable becomes questionable: fitnesses of recessive lethal heterozygotes are typically slightly below maximal (Simmons and Crow 1977). The analyses of (Fig. 1 were therefore conducted again but assigning a viability factor of 0.98 to all *Dd* genotypes. (*A Dd* produced by selfing had fitness 0.9*8cr.)* Although quantitative effects of this fitness adjustment were noted, they were slight, and a parallel figure to that of 1 was effectively indistinguishable from the original (not shown).

**Figure 1.**
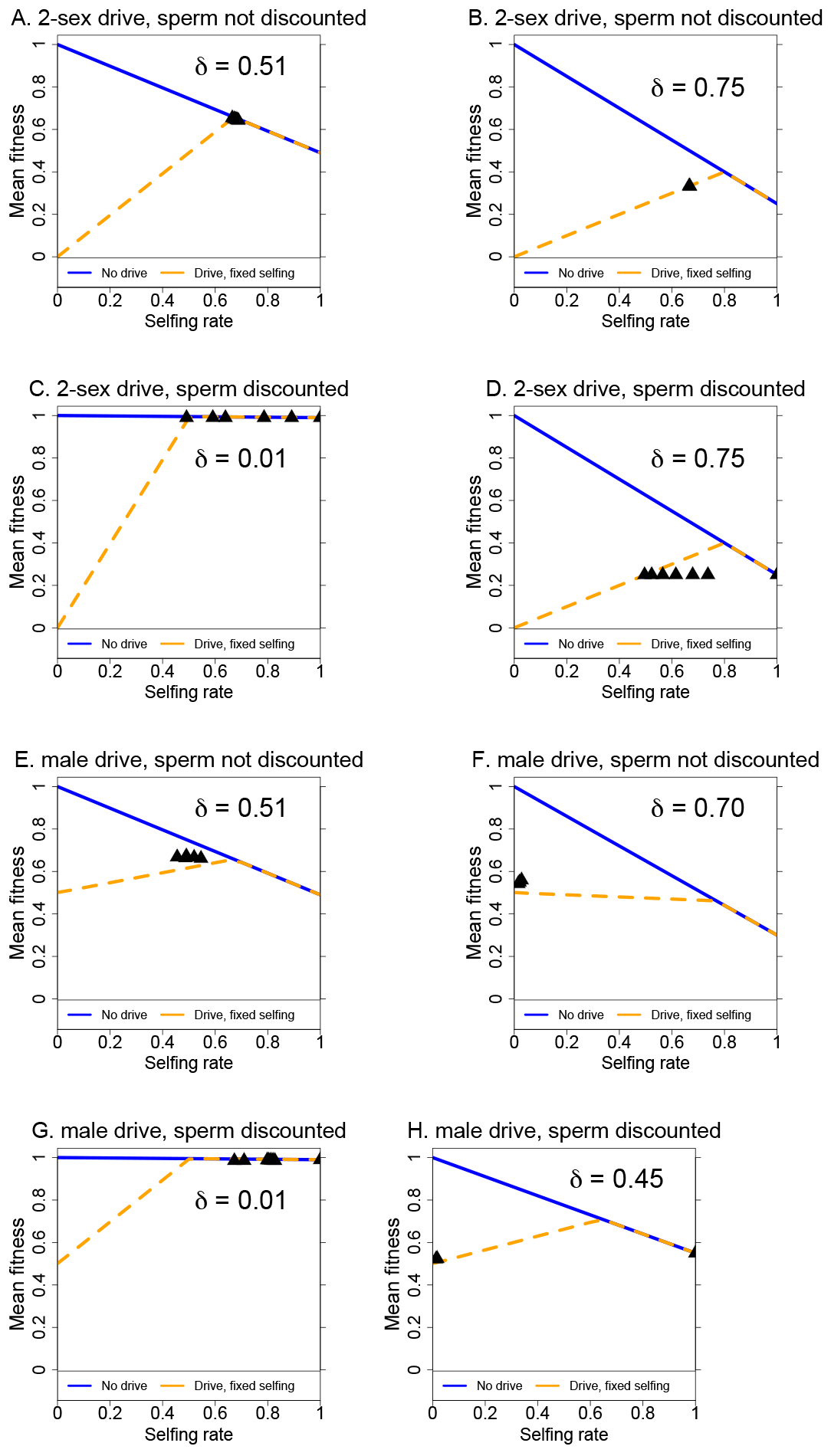
Outcomes of numerical trials for the four models, each evaluated at two different inbreeding depression values *(δ),* as shown. The blue line is mean fitness in the absence of the HEG (drive allele D is absent); the selfing rate is given on the lower axis. Mean fitness declines because higher levels of selfing impose inbreeding depression on more individuals. The yellow, dashed line is equilibrium mean fitness when D has evolved to equilibrium, under a uniform selfing rate given on the lower axis. D is lost for all points on the yellow curve that coincide with the blue line. The black triangles represent equilibria when selfing was allowed to evolve in the presence of the HEG, showing outcomes at or near the highest mean fitness observed across hundreds of trials with different selfing genotypes (selfing rate is the population average selfing rate at equilibrium). For each *δ* value, four sets of initial genotype frequencies were analyzed at each of the 120 combinations of si and s2 values incremented by 0.1 across [0,1] (si = s2 = 0.0 was omitted). Initial genoytpe frequency sets were (0.49, 0.49, 0.01, 0.01), (0.93, 0.05, 0.01, 0.01), (0.05, 0.93, 0.01, 0.01), (0.25, 0.25, 0.25, 0.25) corresponding to (*aadd, aaDd, Aadd, AaDd*); initial frequencies of *AAdd* and *AADd* were 0.

**Figure 2.**
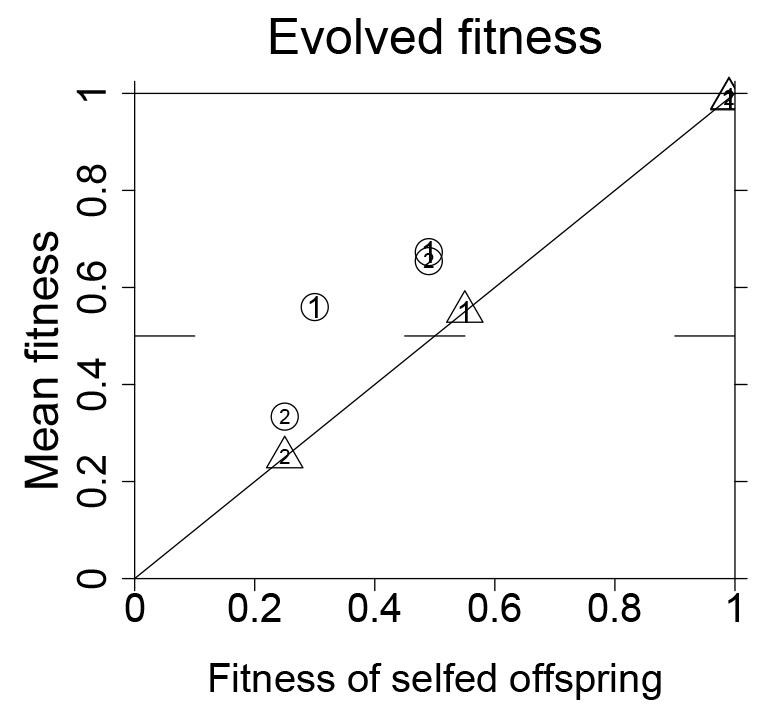
Maximal mean fitness observed in the different models (from Fig. 1) plotted as a function of the fitness of selfed offspring *(σ* = 1 *— δ).* Triangles indicate models with sperm discounting, circles are for models without sperm discounting. A ‘*Y* or ‘*T* is shown within each symbol, indicating the number of sexes exhibiting drive of the D allele. The plot shows that fitness of selfed offspring correlates essentially perfectly with the maximum evolved fitness in the sperm discounting model. Evolved fitness is somewhat higher than *σ* when sperm are not discounted, at least for intermediate values of selfed offspring fitness.

### Invasion and long term fitness: a case study of the male drive, sperm compensated model

The results in Fig. 1 and Fig. 2 do not give insight to which allelic selfing rates were favored. This evolutionary property was explored in detail for the two cases of Fig. 1E and F, the model of male drive with no sperm discounting This case is interesting because mean fitness with the HEG alone evolves to 0.5, and the evolution of selfing is correspondingly prevented at high magnitudes of inbreeding depression.

The equations for this model were linearized for an invasion analysis of the selfing allele A. (Linearization assumed the equilibrium at which the HEG had gone to fixation with *aabb* absent, *aaDd* fixed.) For the genotype frequency vector (*Aadd*, *A Add, AaDd*, *AADd)^f^,* the transition matrix is with leading eigenvalue

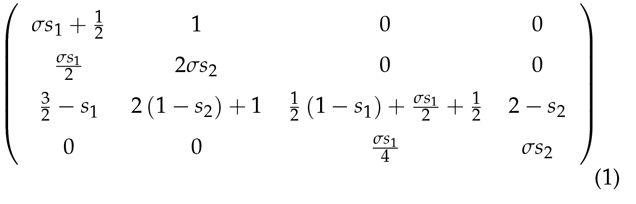

**Figure 3.**
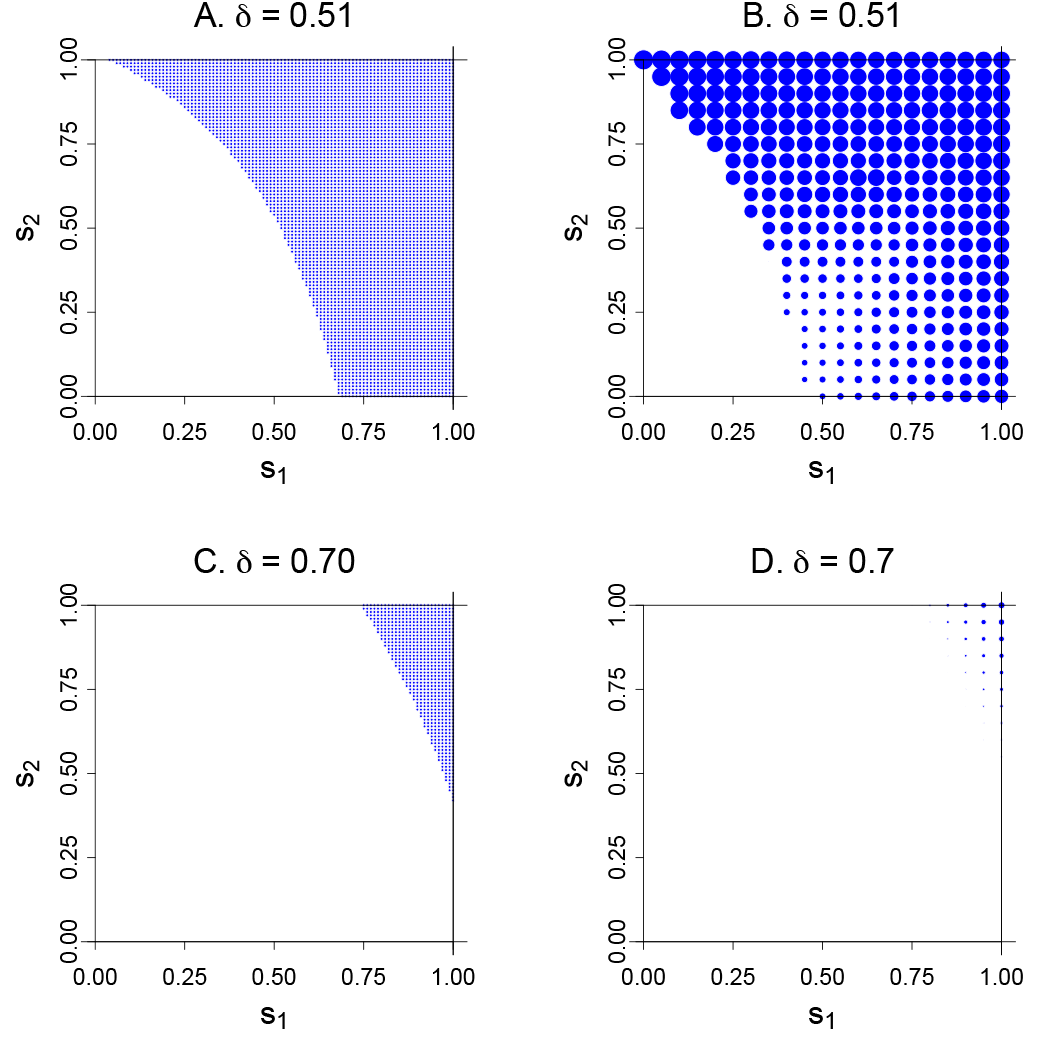
Invasion and equilibrium as a function of genotype self-ing rates (si, S2) for the two models in Fig. 1E, F. Panels (A) and (C) show in blue the regions of the parameter space for which invasion of the selfing allele occurs, from Equation 1. Panels (B) and (D) show equilibrium fitness attained by the allele, in which the diameter of the blue dot is proportional to the excess in fitness above 0.50. Maximal fitness in (B) is fitness 0.67 at Si = 0, S2 = 1. Maximal fitness in (D) is 0.56 at si = 1, S2 = 1. The analytically-derived invasion conditions of (A) are seen to occupy a smaller range than the simulated conditions in (B). These discrepancies arise from the initial frequencies of allele *A* in the numerical trials being higher than is appropriate for the linearization assumptions. In (B), the region of invasion but low fitness (small points in the low, center part of the panel) arise from the allelic constraints preventing the evolution of high selfing rates and thus preventing evolution of high mean fitness.

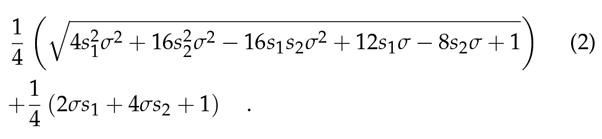

Regions of the (s1, s2) space where this eigenvalue exceeds 1.0 are plotted in blue in Fig. 3A and C (for the values of δ)=)0.51 and 0.70, respectively). Evolved mean fitness values are given in panels (B)and (D); the diameter of each blue dot in (B) and (D) is proportional to the equilibrium fitness. It is evident for both cases, moreso for δ)=)0.7, that evolution of selfing requires that the *A* allele enact high levels of selfing in either or both *Aa* and *AA.* Small levels of selfing do not invade, and as seen in Fig. 3B and D, the highest equilibrium fitnesses occur with s1 and/or s2 at or near 1.0. Selfing will not evolve in this system incrementally.

A similar pattern was evident for the two models analyzed of male drive with sperm discounting (not shown). Invasion of the selfing allele A occurred for a much wider range of (s1, s2) values for δ)=)0.01 than for δ)=)0.45; again, values near (0,0) did not invade.

However, for both models of 2-sex drive, invasion occurred for even the smallest selfing values tested (0.01,0.01). [Initial frequencies in these runs were (0.05, 0.93, 0.01, 0.01, 0, 0) for *(aadd, aaDd, Aadd, AaDd, AAdd* and *AADd).]*

The male-drive models have another interesting property regarding the evolution of selfing. At equilibrium with 100% segregation distortion and all individuals *Dd*, a mutant *A* allele will always arise in the *Dd* background, yielding a *AaDd* genotype. Henceforth, there will be no way for *A* to become paired with the *dd* genotype because all pollen it receives are D. Complete coupling with D ensures that the increased selfing imposes a load on *A* from the formation of *DD* homozygotes due to selfing. Allele *A* cannot escape this load and is selected against relative to allele *a.* This pathology disappears if segregation distortion is less than complete, or if the starting population is not 100% *Dd*, because *Aadd* is then formed.

### Paradoxical loss of drive for some initial conditions

Differences in initial conditions were sometimes found to have a profound effect on outcomes, indicating the presence of multiple stable equilibria for the same parameter values. The cases of most interest, indeed the only cases in which equilibrium fitness was significantly affected, were those in which the drive allele D was lost for some initial conditions but retained for others. These cases were not analyzed exhaustively, merely being discovered and inspected in the course of trials run for Figs. 1 and 2.

Consider the model and *δ* value in Fig. 1A for s1 = S2 = 0.9. In some trajectories, the drive allele D was sometimes lost rapidly, then followed by the more gradual loss of *A*; loss of the *A* allele of course restores full outcrossing, so this population endpoint is the same as that of the initial invasion. In other trials differing only slightly in initial frequencies, the trajectory was similar initially but then continued with damped cycles between increases in D followed by increases in *A*. A pair of runs with differing outcomes is shown in Fig. 4. The frequency of *D* in (B) dropped below 10^-6^ by generation 34 and remained there for nearly 500 generations, so it is feasible that *D* would be lost in a finite population even if it was retained deterministically.

Loss of *D* and *A* conditional on initial frequencies was also observed for several of the runs in Fig. 1C. In Fig. 1D, initial frequency dependence was observed for some outcomes in which *A* fixed and *D* was lost. All of these latter cases had si = 1.0 and S2 < 1, so the final population had an intermediate level of selfing.

In a strictly infinite population behaving deterministically, *D* would not be expected to be lost completely, and at the point that outcrossing was restored to a high level, *D* should rebound, unless its frequency was so low that it took longer to rebound than the trial continued (up to 10^6^ generations in some trials). It is possible that loss of *D* in these numerical trials was a consequence of limited floating point precision. Of course, on biological grounds, a drop to such low frequencies probably does constitute allele extinction. Nevertheless, and in view of these sensitivities to initial conditions, exhaustive analysis (including stochastic simulations), will be warranted for any application.

### Extended evolution of selfing

The models assumed a single, bi-allelic locus for selfing with an initial state of outcrossing (s0 = 0). It is possible that models allowing an extended evolution of selfing would lead to higher selfing rates with a higher mean fitness. Toward this end, further analysis was carried out of some trials in Fig. 1 in which the *A* allele had fixed and left the population with an intermediate level of selfing (*D* also present). The fixed *A* endpoint was then used to seed trials of the 2-locus model in which the evolved selfing rate was the baseline upon which further variation could act. Figs. 1B, D, and E were evaluated: when a wide spectrum of selfing rates were tested against the new baseline, the opportunity for extended evolution often led to yet higher selfing rates and higher fitness above the starting state. Yet the extended evolution did not exceed the highest mean fitness found in the previous trials, although extended evolution often achieved the same highest fitness, or nearly so.

**Figure 4.**
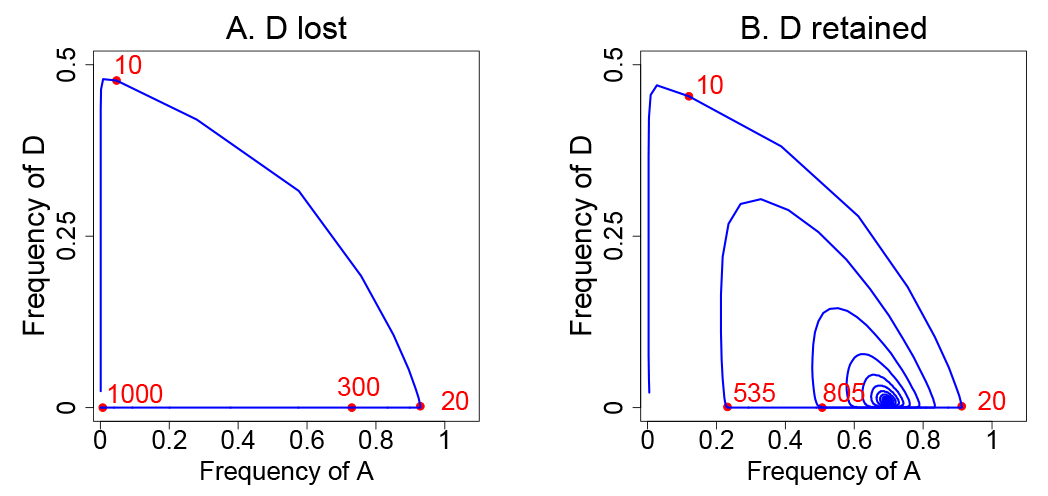
Numerical trajectories of allele frequencies (in the model of 2-sex drive, no sperm discounting) contrasting a case when both *D* and *A* are ultimately lost (A), with a case in which they are both maintained (B). Red dots and associated numbers give the generation number at the point along the trajectory. (A) Initial frequencies of *aadd, aaDd, AaDd, Aadd, AADd* and *AAdd* were 0.95, 0.048, 0.001, 0.001, 0, 0, respectively. The trajectory ends with all individuals being *aadd.* (B) Corresponding initial frequencies were 0.95, 0.040, 0.005, 0.005, 0,0, respectively. In (A), loss of A and D was observed whenever *aadd* was 0.14 or higher (the *aaDd* frequency making up the difference, and other genotype initial frequencies the same). The model used for these dynamics was the one from Fig. 1A. s1 = S2 = 0.9.

Models restricted to two alleles at one selfing locus may also not provide full insight to long term evolutionary dynamics. *A* 3-locus model was created to investigate this possibility (two loci*— A/a* and *B/b —* affected selfing, the *D/d* locus encoded the drive).

The case of Fig. 1H was evaluated, as the mean fitness evolved under the 2-locus model was substantially below the maximum possible in the presence of the HEG. The initial conditions specified selfing rates of 0, 0.9, 0.9 for the *aabb, Aabb,* and *AAbb* genotypes, as led to the modest invasion of selfing in Fig. 1H. The *B* allele forced increases in the lowest level of selfing but maintained 0.9 as the highest level, thus evolution of *B* would have forced the baseline level of selfing above 0. None of those parameter combinations or the others tested resulted in an increase of *B* suggesting that a multi-locus selfing system does not easily evolve to the level of selfing that would maximize mean fitness in the presence of the HEG.

Taken together, these results suggest that the trials for Figs. 1 and 2 captured the highest fitness capable of evolving under selfing. There are clearly constraints on which selfing alleles are favored (e.g., Fig. 3), but the suite of alleles considered may have captured the outcomes with maximal fitness evolution via selfing.

## DISCUSSION

There is much justified excitement about the possibility of employing gene drive systems to limit wild populations of undesirable species (Sinkins and Gould 2006; Gould *et al.* 2008; Burt 2014; Esvelt *et al.* 2014; Unckless et al. 2015). In particular, homing endonuclease genes (HEGs) can now be developed that are recessive lethals but experience close to complete segregation distortion in heterozygotes (Gantz and Bier 2015). Such HEGs can theoretically spread to fixation (extinction) or to the point that the entire viable population is heterozygous (Prout 1953; Bruck 1957; Lewontin 1958), with a major reduction in mean fitness. Evolution of resistance to the HEG, or to its effects, becomes highly relevant in understanding the possible limitations of these engineered systems.

The models analyzed here were of an HEG evolving in a hermaphrodite population that could evolve selfing. There may well eventually be direct applications of these models in weed control (plants being the obvious target species with the highest levels of hermaphroditism), and the frequently observed evolution of selfing in plants (Wright *et al.* 2013) is a caution that such HEG implementations may have limited impact. Yet the results of this study presumably have ramifications for species with males and females, where the most immediate uses of a lethal HEG (and other genetic methods of control) are entertained-insects that transmit disease and destroy crops (Gould *et al.* 2008; Gould 2008). Generality of the present results is in fact suggested by several previous studies on the evolution of mating system evolution in response to a selfish element (see ‘Evolution of mating systems’ below).

This study indicates that the final frequency of a recessive lethal enjoying complete segregation distortion can be reduced by the evolution of inbreeding. There is a consequent increase in mean fitness above that which evolves in the absence of selfing. One important result is that the fitness mitigation achieved by selfing is limited largely by the magnitude of inbreeding depression. In models assuming sperm discounting, selfing enabled mean fitness to avoid the low value expected from an uncontrolled HEG, but mean fitness at equilibrium was the same as that of a selfed offspring *(σ* = 1 — δ). Recovery was somewhat higher in the models with no sperm discounting (Fig. 2).

Although the deleterious population consequences of the HEG could be partly mitigated by the evolution of selfing, it was also true - in male-drive models - that selfing was favored only if the selfing allele enacted a sufficiently large degree of selfing. Alleles with low levels of selfing could not necessarily invade when alleles with high levels could. A conjecture is that an allele needs to attain high levels of selfing to purge the drive allele (which is a recessive lethal) and thereby escape the load that is otherwise associated with inbreeding.

The ultimate fate of the drive allele was sometimes found to depend on initial gene frequencies. For some initial frequencies, the drive allele could invade but was then lost, whereas slight modifications of the initial frequencies led to maintenance of the drive allele. For the sets of initial frequencies tested, long term maintenance of the drive allele was the common outcome (except when complete selfing evolved throughout the population). Yet the mere existence of dynamics in which the HEG invades and is then lost indicates that an extensive analysis of the full parameter space and initial conditions will be needed for any application.

### Precedents in the evolution of mating systems

Evolution of inbreeding in response to a lethal HEG is one of now several examples of mating/genetic system evolution in response to a selfish allele (or alternatively, an altruistic allele). In some cases, it has merely been suggested that inbreeding limits the impact of a selfish element (e.g., Hamilton 1967; Burt 2003). In one study, however, inbreeding was shown to evolve as a response to altruism (Breden and Wade 1991): the allele for increased inbreeding became coupled with the allele for altruism, with inbreeding reinforcing the benefit of altruism by increasing the number of altruists interacting in family units. A few studies have shown that the number of sires - polyandry or monogamy - evolves in response to the presence of a selfish element (Haig and Bergstrom 1995; Champion de Crespigny et al. 2008) or in response to altruism (Peck and Feldman 1988); the number of sires affects relatedness within families and thus affects the competition between selfish and non-selfish alleles. In a system of yet higher complexity, Lande and Wilkinson (1999) found that a sex-linked segregation distorter could be partially or wholly suppressed by female preference of a sex-linked male trait if that trait was tightly linked to the segregation distorter. Brandvain and Coop (2012) found that segregation distorters in female meiosis can select changes in the female recombination rate.

### Limitations of the models

***Other forms of inbreeding*** With many proposed applications of gene drive systems to disease vectors and crop pests (Burt 2003; Gould et al. 2008; Gould 2008), the evolution of inbreeding as a possible ‘escape’ from a lethal HEG is especially relevant to insects. An obvious extension of the work here is to consider the evolution of inbreeding in species with separate sexes, the most straightforward form of inbreeding being sib mating. The evolution of selfing in response to a lethal HEG is likely to generalize to the evolution of sib mating, but one potentially critical difference is that selfing more easily achieves high inbreeding coefficients than does sib mating. As suggested by R. Lande (pers. commun.), the fact that selection favored selfing alleles in our models only if they enacted moderate to high rates of selfing (e.g., Fig. 3) may indicate that the evolution of sib mating will face even greater constraints than does the evolution of selfing. Such a result would raise hope that the evolution of selfing is less likely to thwart an HEG in insects than in selfing species. However, the restrictions against evolution of low selfing rates were observed only for the male-drive models, so it is not clear whether the evolution sib mating would be constrained with 2-sex drive.

***Evolution of inbreeding depression*** The models assumed that inbreeding depression (δ) was fixed throughout the evolutionary process. In contrast, inbreeding depression is known to evolve and can be at least partly overcome when inbreeding is imposed on formerly outcrossing populations (Charlesworth and Willis 2009). In the results here, the magnitude of inbreeding depression played a critical role in invasion and ultimate recovery of population fitness (e.g., Fig. 2). Any purging of inbreeding depression would not necessarily be relevant to invasion conditions, but it would improve the recovery after invasion beyond that seen in models with fixed δ. An important extension of these models is thus to incorporate the evolution of inbreeding depression with the evolution of inbreeding.

Recent work points toward ways in which dynamic inbreeding depression might be accommodated. A consensus is emerging that inbreeding depression is due largely to deleterious mutations, but there are two important classes of deleterious mutations that contribute and have different consequences for purging: weakly deleterious mutations with large additive effects and strongly deleterious mutations such as lethals whose effects are largely recessive (Charlesworth and Willis 2009; Porcher and Lande 2005a, 2013). The weakly deleterious mutations are abundant, whereas the strongly deleterious mutations are much less common. In the short term, purging is chiefly from loss of the strongly deleterious class (Porcher and Lande 2005a, 2013).

An useful extension of the model is to expand the biological ramifications of inbreeding depression to include reproductive compensation for inviable zygotes. In species with post-zygotic parental investment in offspring, reproductive compensation boosts offspring number by replacing inviable genotypes with viable ones (e.g., Porcher and Lande 2005b; Harder et al. 2008). The ramifications of this could be to facilitate the evolution of drive by supplementing *Dd* offspring to replace inviable *DD*.

***Implications for population suppression*** The models analyzed here are of population genetics evolution. Yet interest in applying gene drive systems is often for population control - to limit numbers of adults. Unfortunately, across diverse ecological settings, there is no straightforward relationship between mean fitness and adult population size, (the one exception being that a mean fitness of 0 ensures extinction).

Of particular relevance to population suppression by a recessive lethal HEG is that the lethality will typically operate at the zygotic stage. A reduction in zygotes will be especially prone to be overcome by many types of density-dependent regulation, whereby the number of adults is much less reduced than is the number of zygotes (Foster and Whitten 1974). In the long history of work on the sterile insect technique - the goal of which is to suppress pest populations by artificially introducing sterile males or females - species-specific ecology often underlies the difference between success and failure (Pal and Whitten 1974; Ito and Yamamura 2005; Klassen and Curtis 2005; Lance and McInnis 2005). It is thus to be expected that the impact of gene drive system to suppress a target species will also be ecology dependent, unless of course it can destroy all progeny.

### Future

The empirical feasibility of evolving high levels of inbreeding during assault by a recessive lethal HEG remains to be seen. We have essentially no field experience with gene drive systems. There is, however, over half a century of experience with various forms of sterile insect applications (Bushland *et al.* 1955; Whitten 1971; Dyck *et al.* 2005). Sterile insect techniques almost universally rely on the release of lab-reared insects that, when mated with wild insects, cause death or sterility of the progeny (Sinkins and Gould 2006). (Those lab-reared insects may be irradiated or mutated in other ways or may carry chromosomal rearrangements that are incompatible with the wild population.) The assault from sterilizing, lab-reared insects should also favor inbreeding as one of several mechanisms that avoid matings that produce sterile progeny. Yet assortative mating by wild populations subjected to the sterile insect technique has rarely been reported, despite many applications and successes (summarized in Bull 2015). In this comparison, it may be important that the sterile insect technique relies on inundation of the wild population with lab-reared insects, the wild individuals becoming increasingly overwhelmed with sterility-inducing matings as the population declines. Gene drive systems may have the opposite effect, encouraging *de facto* consanguinity as the population density declines. Nonetheless, there are many reasons to be hopeful that gene drive systems will be able to achieve long-standing population control in at least some species.

## ACKNOWLEDGMENTS

R. Lande read early drafts, provided extensive comments and suggested analyses that greatly improved the study. S. Wright and two anonymous reviewers also helped, especially with literature and general recommendations. Mathematica 10.4.0.0 was used in some analyses. Supported by NIH GM 57756.

## APPENDIX

The basic recursion equations are common to all models. Assuming discrete, non-overlapping generations:

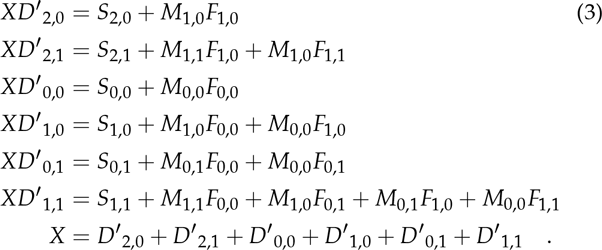

Here, D;,j represents a diploid adult, with *i A* alleles and *j D* alleles. Diploids may have 0,1 or 2 A alleles but only 0 or 1 D alleles because D is a recessive lethal. Mk,_n_ (Fk,_n_) represents outcrossed male (female) gametes with *k A* alleles and *n D* alleles. S_i,j_ represents diploid offspring by selfing, with similar subscripting as for *D_i j_*. Recursion equations for the gametes and selfed offspring are specific to each model, as follows. It is assumed that all ova are fertilized, and sperm frequencies are normalized. Although the body of the paper restricts the analyses to complete drive, these equations use an unsubscripted D to represent the fraction of gametes of a D heterozygote carrying the D allele, facilitating the derivation. Selfing rates are so for *aa,* si for *Aa*, and S2 for *AA*, regardless of the other locus.

**Male-female drive, sperm discounted**

***Sperm***

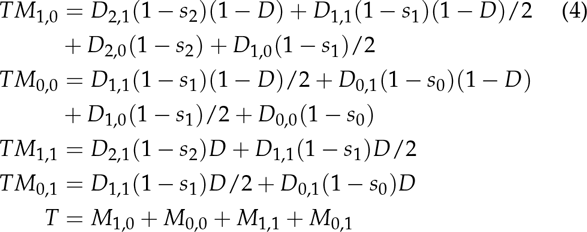

***Ova***

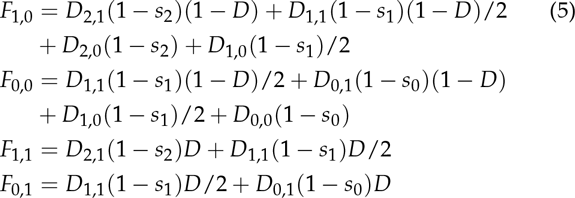

***Selfed***

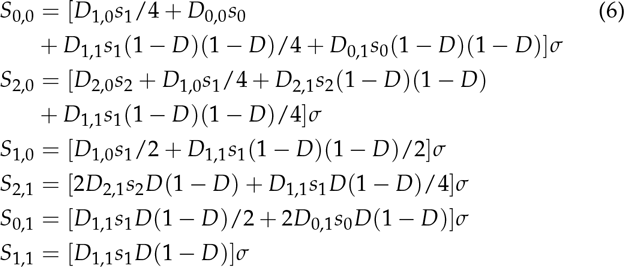

**Male-female drive, sperm compensated**

***Sperm***

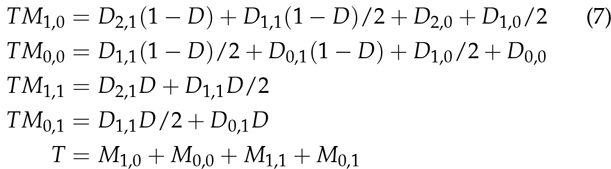

***Ova***

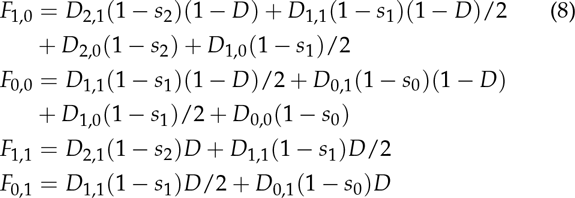

***Selfed***

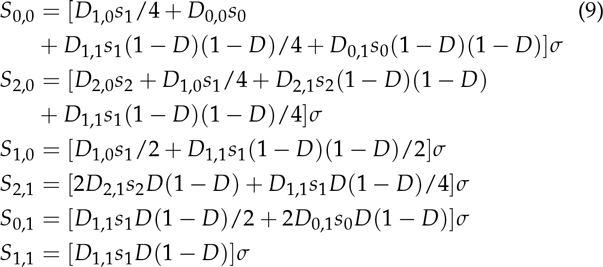

**Male drive, sperm compensated**

***Sperm***

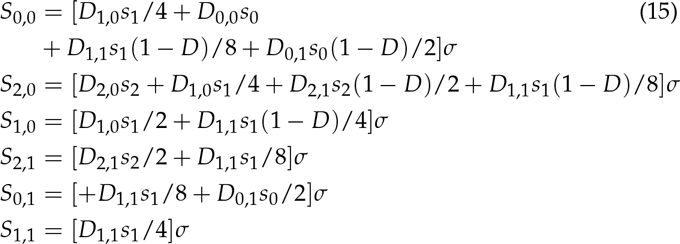

***Ova***

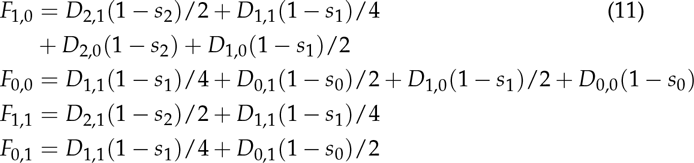

***Selfed***

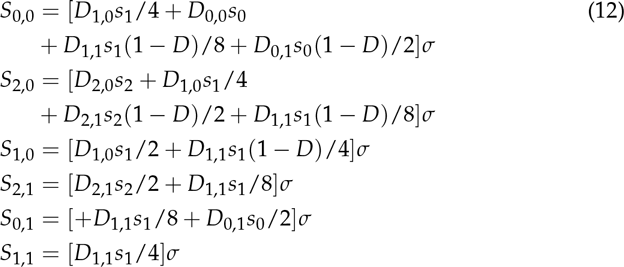

**Male drive, sperm discounted**

***Sperm***

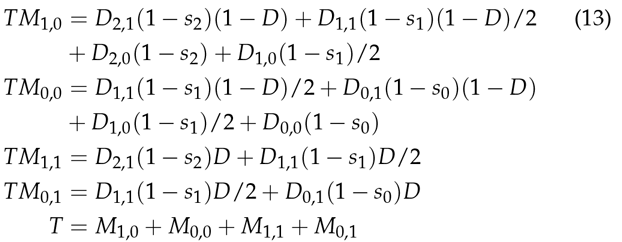

***Ova***

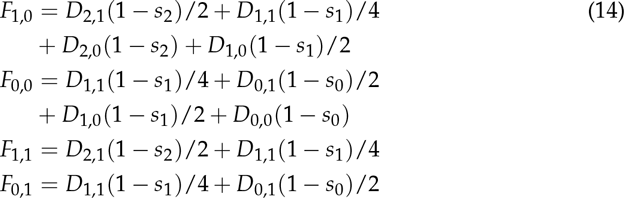

***Selfed***

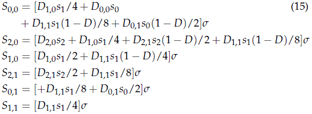

